# A Deep Learning Model for Accurate Segmentation of the *Drosophila melanogaster* Brain from Micro-CT Imaging

**DOI:** 10.1101/2024.12.30.630782

**Authors:** Jacob. F. McDaniel, Mike Marsh, Todd Schoborg

## Abstract

The use of microcomputed tomography (Micro-CT) for imaging biological samples has burgeoned in the past decade, due to increased access to scanning platforms, ease of operation, isotropic three-dimensional image information, and the ability to derive accurate quantitative data. However, manual data analysis of Micro-CT images can be laborious and time intensive. Deep learning offers the ability to streamline this process, but historically has included caveats—namely, the need for a large amount of training data, which is often limited in many Micro-CT studies. Here we show that accurate deep learning models can be trained using only 1-3 Micro-CT images of the adult *Drosophila melanogaster* brain using Dragonfly’s pre-trained neural networks and minimal user knowledge. We further demonstrate the power of our model by showing that it can accurately segment the brain across different tissue contrast stains, scanner models, and genotypes. Finally, we show how the model can assist in identifying morphological similarities and differences between mutants based on volumetric quantification, facilitating a rapid assessment of novel phenotypes. Our models are freely available and can be refined based on individual user needs.

**Summary:** Micro-CT data can be automatically segmented and quantified using a deep learning model trained on as few as 3 samples, facilitating rapid comparison of developmental phenotypes.

## Introduction

X-Ray based imaging methods, such as computed tomography (CAT) and its higher resolution counterpart, Microcomputed tomography (Micro-CT), are advantageous for their nondestructive nature and ability to generate three dimensional (3D) images of a wide variety of materials. As a result, they have become common in a variety of fields, including materials science, geology, biology, and medicine (Buffiere et al., 2010; Carlson, 2006; Hounsfield, 1980; Schoborg et al., 2019). With the proliferation of improved and standardized tissue staining techniques, Micro-CT has established a niche in developmental biology as an effective imaging method for analyzing soft tissues as well as more dense structures such as bone (Clark and Badea, 2014; du Plessis et al., 2017; Keklikoglou et al., 2021; Metscher, 2009; Singhal et al., 2013).

Despite these advances, Micro-CT remains an underutilized imaging method due in part to the laborious process of image analysis and quantification, which is essential for realizing the true power of this imaging technology for scientific investigation. Manual whole organ segmentation from entire specimens are especially labor intensive. Although many automated and semi-automated segmentation methods have been devised to address this issue, many of them also require extensive manual input to accurately segment structures of interest, thus limiting their feasibility (Chai et al., 2023; McGrath et al., 2020; Yushkevich et al., 2006). Furthermore, deep learning for image segmentation, while powerful, has historically been inaccessible to many labs due to the volume of data required (>thousands of training datasets) and the required expertise in computational science (Lee et al., 2022; Sapoval et al., 2022).

Recently, many image analysis software platforms have implemented deep learning solutions to help end-users overcome these limitations in image processing and analysis. For example, Dragonfly has developed several innovations to automate this task, including a series of pre-built and pre-trained neural networks. Other improvements include a deep learning wizard to improve accessibility and lower the learning curve for novice users (Makovetsky et al., 2018), data augmentation techniques to reduce sample size and improve scalability (Provencher et al., 2019), as well as improved deep learning models specifically designed to segment by shape (Badran et al., 2020). Models can also be recycled and repurposed by other users for their own image analysis needs. These innovations, coupled with Dragonfly’s free academic license, prompted us to evaluate if it could be used to automate the laborious effort of manually segmenting *Drosophila melanogaster* organs from Micro-CT scans (Schoborg et al., 2019).

Here we demonstrate the ease and feasibility by which deep learning tools can accelerate the analysis of complex 3D Micro-CT datasets. We used the *Drosophila melanogaster* brain and visual system as our targeted structures and built a series of models that can segment with a high degree of accuracy (>98% for total volume, >95% for individual regions in non-mutant flies) and precision compared to manual methods. The simplest models, consisting of just 1-3 training datasets, are suitable for images that use the same sample parameters, such as stain type, Micro-CT scanner model, and genotype. Using these models as a baseline, we then built into them the ability to handle different imaging and sample parameters. In each case, we found that providing 6 samples of training data is generally sufficient when adding a new sample parameter. For example, we show that a single deep learning model trained with 12 datasets can accurately predict fly brain regions from images taken on two different Micro-CT scanner models, while a model with 43 datasets was sufficient to segment images taken with 2 different scanners, stains, and mutant genotypes.

Together, our results show that Dragonfly’s Deep Learning tools consistently create models capable of accurate, reproducible organ segmentation and volumetric quantification, with a 10-fold increase in speed compared to manual segmentations. With our framework of pretrained models, other labs can retrain and refine them for their own needs using minimal training data, thus lowering the burden of Micro-CT imaging and its associated analysis.

## Results and Discussion

We first set out to discover the minimum number of training images required to create a deep learning model capable of segmenting an adult *Drosophila* brain imaged by Micro-CT (Fig. 1A,B). The *Drosophila* brain consists of three large structures: a central brain sandwiched between two optic lobes, which can be resolved by the 3-5 μm image resolution capable of many Micro-CT scanners (Ito et al., 2014). The lamina, located distally to each optic lobe, helps relay visual information from the retina to the optic lobes (Fig. 1C).

**Figure 1.**
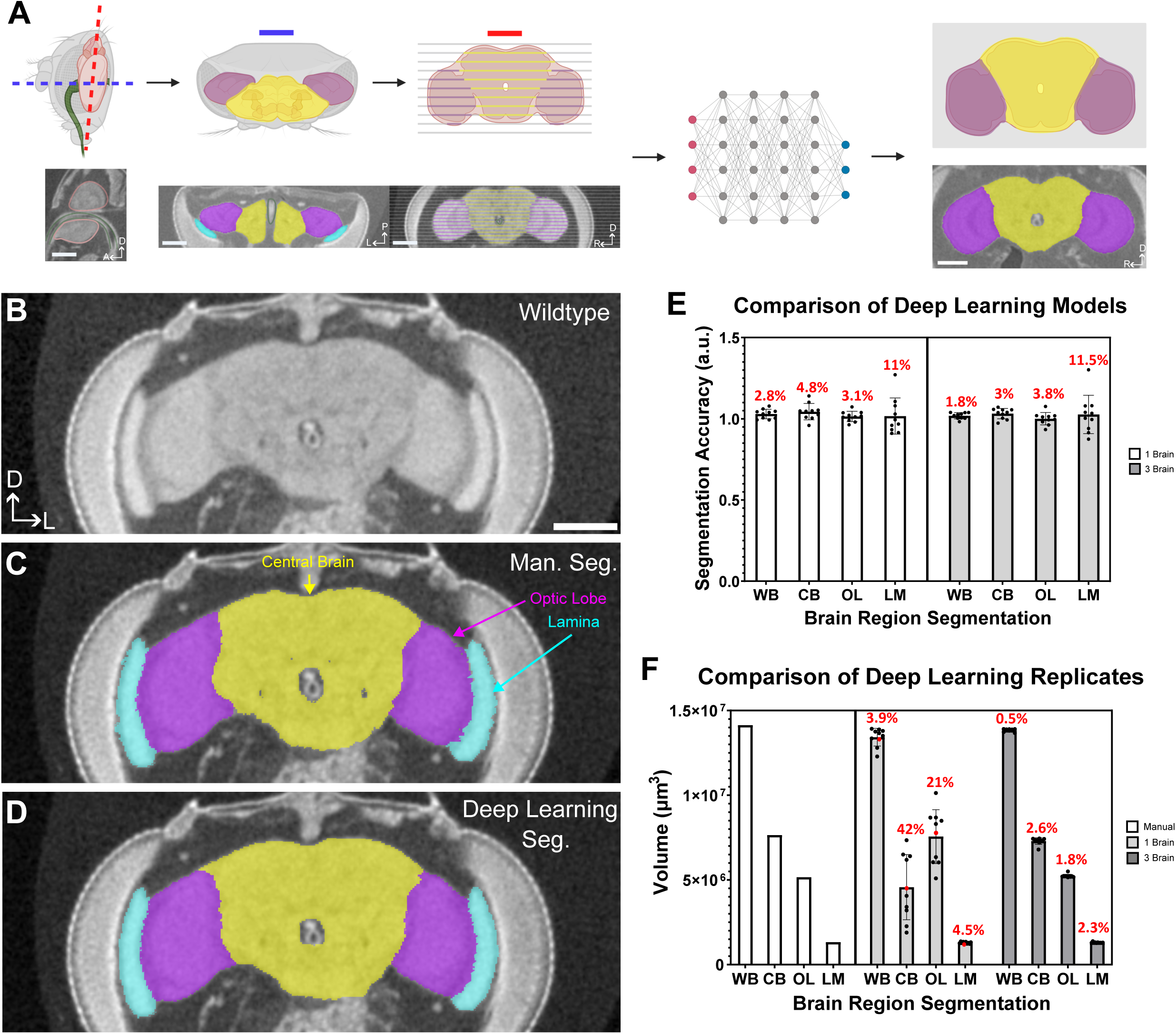
Accurate fly brain segmentation models using deep learning can be generated from only one to three Micro-CT images. (A) Schematic overview of Dragonfly’s deep learning workflow. Models were trained and segmented along the transverse plane (blue line), while most images are shown along the coronal plane (red line). See Methods for details. (B) Representative section of a wildtype adult *Drosophila melanogaster* head case imaged on the Zeiss Xradia Versa 620. (C) Representative manual segmentation of the brain and associated lamina neuropil. Yellow = Central Brain, Purple = Optic Lobes, Cyan = Lamina. (D) Representative deep learning segmentation of the same brain, using the 3 brain model. (E) Comparison of deep learning model accuracy between the one and three brain models. Segmentation accuracy is the volume of a manual segmentation divided by the volume of the specified deep learning model (n=10 brains). Whole Brain (WB), Central Brain (CB), Optic Lobe (OL), and Lamina (LM). (F) Reproducibility of the one and three brain models (see methods). Model precision was calculated using the coefficient of variation (CV), shown in red % above each bar in (E) and (F). n≥10 brains, Welch’s t-test. ns, P>0.05; *P≤0.05; **P≤0.01; ***P≤0.001; ****P≤0.0001. Error bars represent standard deviation. Anterior (A), Posterior (P), Dorsal (D), Left (L), Right (R). Scale bars are 100 μm.

We found that just a single training image consisting of 20% ground truth (every 5^th^ slice manually segmented, Fig 1A) was sufficient to create a single model (1 Brain) that was able to segment the whole brain (WB), central brain (CB), optic lobes (OL), and lamina (LM) with >98% accuracy and a high degree of precision (Coefficient of Variation (CV) 2.8-11%) from 10 different brains imaged with the same conditions (Fig. 1D,E). We define ‘accuracy’ as the ratio of the normalized brain volumetric measurement (μm^3^/thorax width, μm^2^) (Schoborg et al., 2019) from the model segmentation to the normalized volumetric measurement from the manual segmentation of the same brain, which we take to be the true/accepted volume value.

The performance of our 1 Brain model was on par with a second model that we built, which consisted of 20% ground truth each from three different brain images (3 Brain). The accuracy and precision of both models were comparable (>98% Accuracy, 1.8-11.5% CV; Fig. 1E, Video 1 and 2). We conclude that high performing models can be trained from a minimal amount of ground truth and computational knowledge, thus limiting the burden to investigators.

We next wanted to address the reproducibility of our models. We therefore trained ten different 1- and 3-Brain models using the same 20% ground truth for each training session, then compared each model’s accuracy and precision in predicting brain volumes from the same 10 brain images (Fig. 1F). We observed a larger variation (4-42% CV) in the ability of each one brain model to accurately predict volumes, particularly for the central brain and optic lobes (Fig. 1F). However, the 3 brain models were much more precise (0.5%-2.6% CV) with a higher degree of accuracy (Fig. 1F). Thus, while accurate segmentation models can be obtained from just a single training image, incorporating three images for training greatly improves the reproducibility of the models. It is also important to validate these models against a manually segmented image to ensure the model is as accurate as possible prior to use.

While these models performed well given the consistent imaging conditions (same Micro-CT scanner, iodine stain for tissue contrast, wildtype brain morphology), we next sought to determine how altering these variables would affect the amount of training data required to generate a highly robust segmentation model. First, we addressed if a segmentation model could be trained to accurately detect brain regions from images taken on different Micro-CT scanners— the Zeiss Xradia Versa 610 (Fig. 1) and the Bruker Skyscan 1172 (Fig. 2). When we attempted to segment data imaged on the Skyscan (Fig. 2A) using the 3-brain model that was trained on the Versa images, we found that the model struggled to accurately segment each brain region (Fig. 2C) compared to manual segmentation (Fig. 2B) and was highly variable (29-87% CV, Fig. 2E). To resolve this, we iteratively added additional training data to a new model until we had 6 samples from each scanner. Using six brains from each scanner (12 total) as training data allowed us to generate a highly accurate and precise (5-15% CV) multivariable 12 brain model that could predict brain regions from images taken with either scanner. With this quantity of training data, the model was able to consistently segment samples (Fig. 2D) and achieve >98% accuracy on the full brain, and >97% accuracy for the central brain and optic lobes individually, regardless of the scanner used for imaging (Fig.2E).

**Figure 2.**
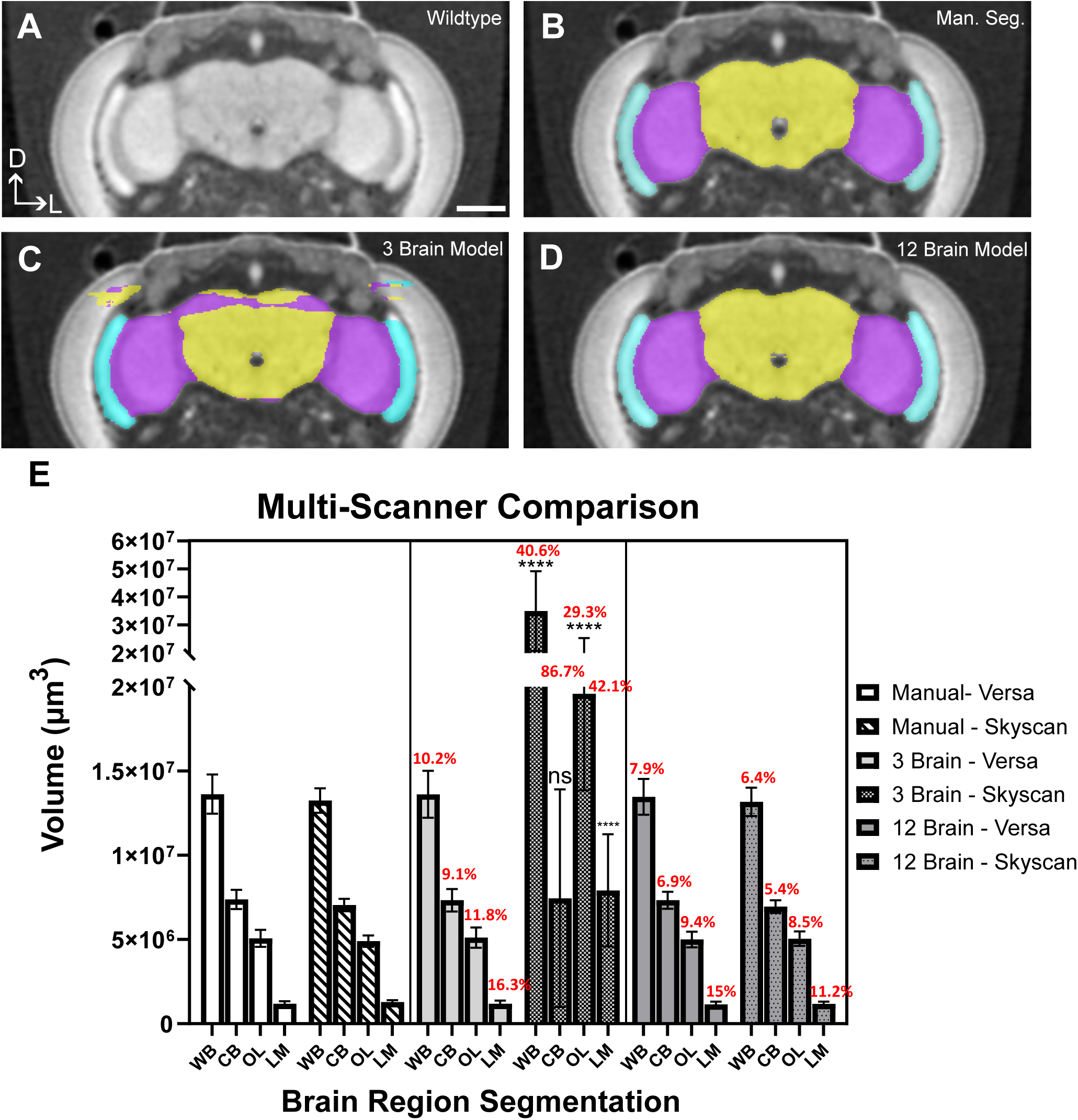
A single deep learning model can be trained to identify fly brains imaged on different Micro-CT scanners. (A) Representative section of a wildtype adult *Drosophila melanogaster* head case imaged on the Bruker Skyscan 1172. (B) Representative manual segmentation of the brain and associated lamina neuropil. Yellow = Central Brain, Purple = Optic Lobes, Cyan = Lamina. (C) Representative deep learning segmentation of a Skyscan image segmented using the 3 brain model trained only on Zeiss Xradia Versa images. (D) Representative deep learning segmentation of a Skyscan image segmented using a 12 brain mixed model that included 6 Skyscan and 6 Versa images in the training dataset. (E) Volumetric quantification of segmentations from 3 brain and 12 brain mixed models. Whole Brain (WB), Central Brain (CB), Optic Lobe (OL), and Lamina (LM). Model precision was calculated using the coefficient of variation (CV), shown in red % above each bar. n≥10 brains, Welch’s t-test. ns, P>0.05; *P≤0.05; **P≤0.01; ***P≤0.001; ****P≤0.0001. Error bars represent standard deviation. Dorsal (D), Left (L). Scale bars are 100 μm.

We next tested whether changing multiple imaging variables (scanner type, staining protocols, and developmental phenotypes) could be incorporated into a comprehensive mixed model that could accurately and precisely segment the brain. We used animals carrying mutations in the microcephaly gene *abnormal spindle* (*asp*) as our developmental phenotype (Schoborg et al., 2015). *Asp* mutant brains display severe but highly variable morphology defects compared to wildtype brains, particularly in the optic lobe neuropils and lamina (Mannino et al., 2023; Schoborg et al., 2019).

To effectively train this model, we started with 6 samples of training data for our typical imaging condition and 6 samples for each of our major imaging variables (scanner, stain, mutant), as that was sufficient for the two separate imaging conditions in our 12 brain model. We also included 3 samples of each additional possible combination of imaging conditions (e.g., *asp* mutants stained with PTA, imaged on the Skyscan), totaling 36 pieces of training data. However, this model struggled with *asp* mutants stained with iodine, so we continued to add training data until we were satisfied with the model’s performance, finally arriving at 43 samples of training data. In total, this comprehensive model had 43 samples in the training dataset. This model is >98% accurate for total brain volume, central brain, and optic lobe volume for samples with similar imaging conditions and developmental phenotypes as the training data (Fig. 3).

**Figure 3.**
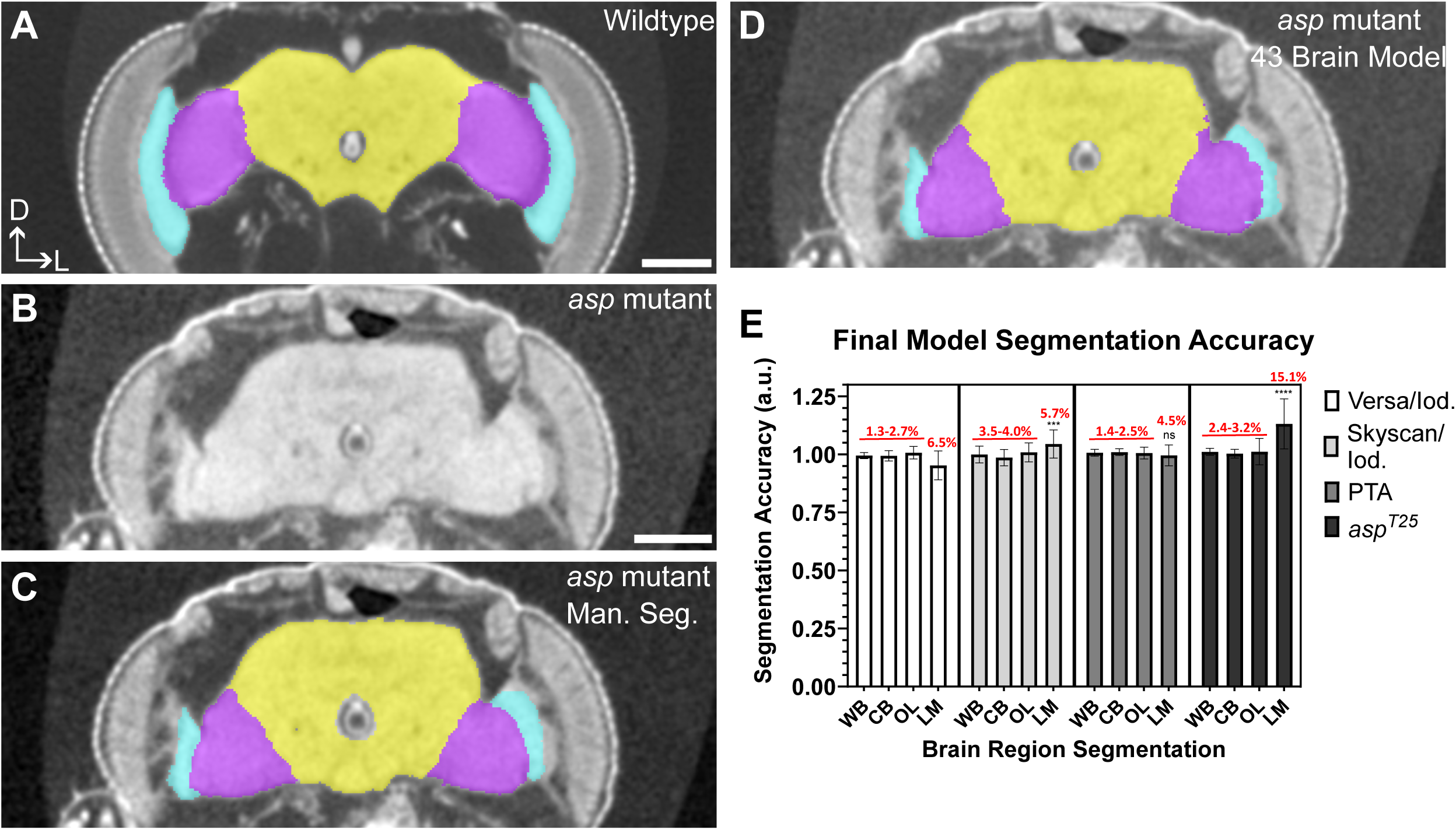
A robust comprehensive model that accounts for different scanner models, contrast agents, and tissue phenotypes can be trained to identify fly brains using limited training data. (A) Deep learning segmentation of a phosphotungstic acid (PTA)-stained *Drosophila* brain imaged on the Versa. Yellow = Central Brain, Purple = Optic Lobes, Cyan = Lamina. (B) Representative image of an *asp* mutant headcase with defective neuropil architecture and microcephaly, stained with iodine and imaged on the Versa. (C) Manual segmentation of the *asp* mutant brain shown in (B). (D) Deep learning segmentation of the same *asp* mutant using the 43 brain comprehensive model. (E) Comparison of final model accuracy for each imaging condition. Whole Brain (WB), Central Brain (CB), Optic Lobe (OL), and Lamina (LM). Model precision was calculated using the coefficient of variation (CV), shown in red % above each bar. n≥10 brains, Welch’s t-test. ns, P>0.05; *P≤0.05; **P≤0.01; ***P≤0.001; ****P≤0.0001. Error bars represent standard deviation. Dorsal (D), Left (L). Scale bars are 100 μm.

It also was very precise for the whole brain, central brain, and optic lobe (<4% CV). However, this model was less precise for the lamina (4.5-15% CV), particularly for the *asp* mutants (Fig. 3E). This is perhaps not surprising, considering the severe and highly variable morphological defects in this region(Schoborg et al., 2019). It may also be due to the lamina’s high surface area to volume ratio, which may make the tissue susceptible to variations in iodine or PTA staining. This in turn can influence the manual segmentation process for images used in the training dataset, due to variability in pixel thresholding. Incorporating more training datasets from *asp* mutants would be expected to increase the model’s accuracy and precision in predicting the lamina, although there may be a point of diminishing returns for training models to predict highly variable tissue phenotypes.

With a comprehensive model in hand, our next objective was to determine if this model could accurately segment phenotypically similar mutant developmental phenotypes it had not been explicitly trained on. To evaluate this model, we attempted to segment *ey^D1Da^* mutants using our comprehensive model that had been trained on *asp* mutants. We chose *ey^D1Da^* mutants as they were previously shown to have a milder phenotype than other *ey* mutants, and also showed similar morphological defects in the optic lobes as *asp* mutants (Fig. 4A). *Ey* expression is also significantly downregulated in *asp* mutants (Callaerts et al., 2001; Clements et al., 2009; Mannino et al., 2023; Schoborg et al., 2019).

**Figure 4.**
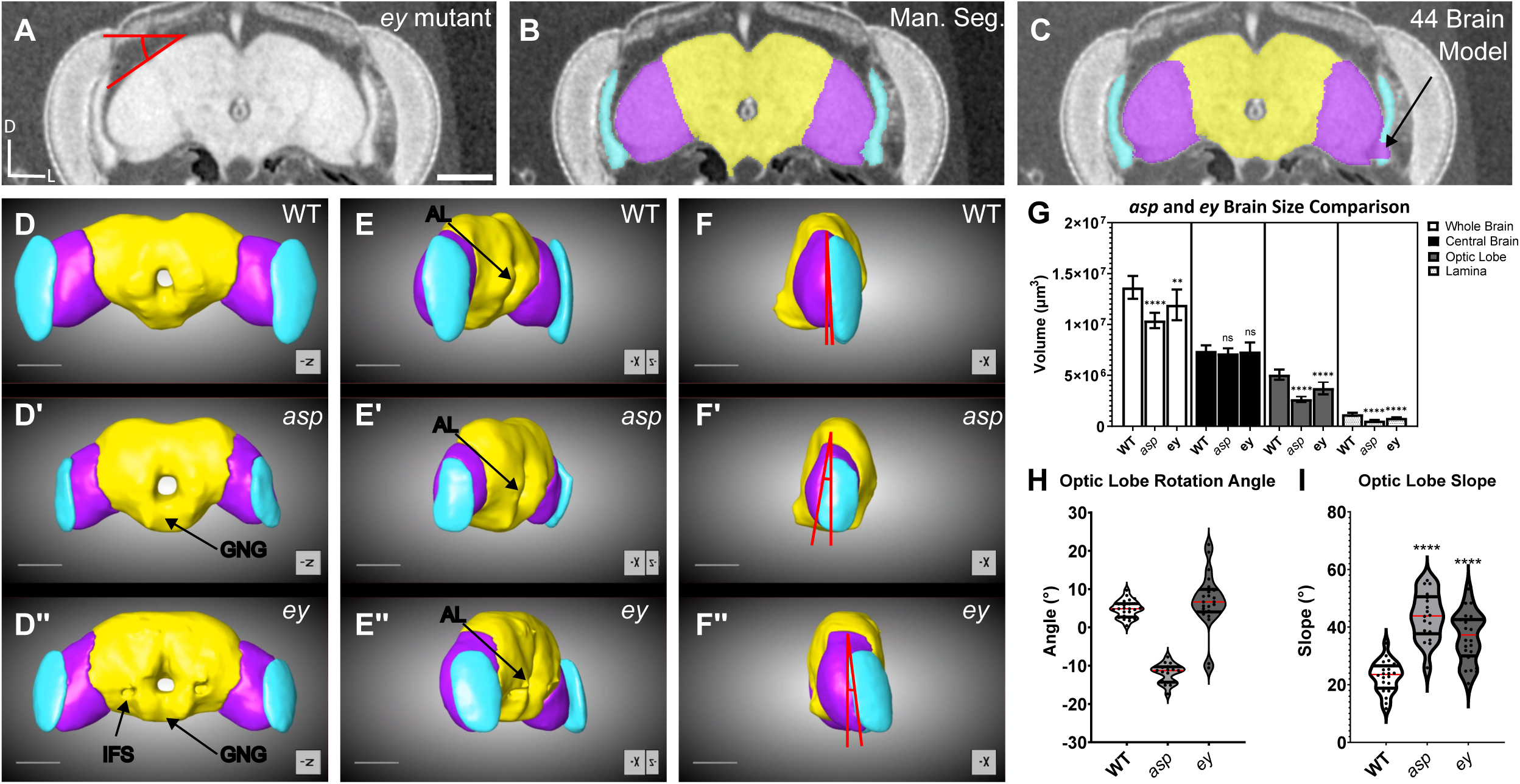
Deep learning segmentation can be used to quickly segment untrained mutants and quantify new phenotypes. (A) Representative *eyeless (ey)* mutant (*ey^D1DA^)*. The angle demonstrates how the optic lobe slope was measured in Fig. 4I. (B) Manual segmentation of *ey* mutant. (C) Deep learning segmentation of *ey* mutant using the 43 brain comprehensive model. Note the categorization of a small portion of the lamina as optic lobe. 3D frontal view of (D) wildtype, (D’) *asp*, and (D’’) *ey* mutants. Note the exposure of the inferior fiber system (IFS) in the *ey* mutant, and reduction in volume of the gnathal ganglion (GNG) in both mutants. 60-degree rotation of the (E) wildtype, (E’) *asp*, and (E”) *ey* brains. Note the reduction in volume of the antennal lobe (AL). 90-degree rotation of the (F) wildtype, (F’) *asp*, and (F*”) ey* brains. Red angle highlights defects in optic lobe neuropil (purple) rotation relative to the central brain (yellow). (G) Quantification of brain volume in wildtype (WT), *asp,* and *ey* mutants. (H) Violin plot showing optic lobe rotation angle, measured as shown in F-F”. Red line denotes the median. (I) Violin plot showing the optic lobe slope, measured as shown in (A). Red line denotes the median. n≥10 brains, Welch’s t-test. ns, P>0.05; *P≤0.05; **P≤0.01; ***P≤0.001; ****P≤0.0001. Error bars represent standard deviation. Dorsal (D), Left (L). Scale bars are 100 μm.

Upon segmentation of *ey* mutants (Fig. 4), we noted that the central brain and other regions that were visually similar to *asp* or wildtype brains were accurately segmented, while areas that were less similar such as the lamina had some imperfections (Fig. 4C). This suggests that while our model is capable of accurately segmenting similar mutant phenotypes without explicit training, results should always be closely evaluated when comparing dissimilar tissue between mutants.

However, we did uncover an unexpected advantage of employing a model on untrained mutant images—its ability to reveal new/overlooked phenotypes. For example, *ey* mutants have not been explicitly reported to have microcephaly, yet our volume assessment using our trained *asp* model clearly showed that these animals, much like *asp* mutants, also have microcephaly, with the *ey* mutant optic lobes showing the greatest reduction in size (Fig. 4G).

Furthermore, a ‘mistake’ in the automated segmentation of untrained mutants can also reveal clues to additional phenotypic defects that may not be obvious to the human eye. For example, a comparison of manual and automatic segmentations revealed a previously unnoticed morphological phenotype in the central brain of some of the more severe *ey* mutants (Fig. 4D-4D”, 4E-4E”, 3/12 animals). This phenotype exposes the inferior fiber system (IFS). In addition, there is a visible reduction in size of the gnathal ganglion (GNG) and antennal lobe (AL) in both mutant brains when compared to wild type. Other well characterized *ey* defects, such as mushroom body size and morphology, were also clearly evident in our analysis(Kurusu et al., 2000). Interestingly, overall central brain volume did not appear to be significantly affected, despite the morphological differences observed (Fig. 4G), which may be due to compensatory overgrowth in other neuropil regions in this part of the brain. This could be revealed by additional AI models that are trained to identify individual fly neuropils, rather than larger brain structures. These ‘high-resolution’ models would also be expected to accelerate the identification of new mutant phenotypes elsewhere in the brain.

In support of this idea, we also observed additional defects in the optic lobe neuropils in both *asp* and *ey* mutants. The fly optic lob consists of only three neuropils (medulla, lobula, and lobula plate), making it a much ‘simpler’ structure than the central brain. In this case, our model not only highlighted a size (volume) phenotype in *asp* and *ey* mutants (Fig. 4G), but it also revealed a unique arrangement of these neuropils compared to wildtype brains.

Upon analyzing the three-dimensional segmented structures of these mutants (Video 3), we noted that the optic lobes and lamina of *asp* (Fig. 4F’) and *ey* (Fig. *4*F”) mutants were ‘rotated’ in different directions compared to wild type brains when viewed from the sagittal perspective. When this rotation was measured and quantified (Fig. 4F’, 4F”, H), *asp* mutants were biased towards a negative angle of rotation, while *ey* mutants showed severe rotation angles in both directions. In addition, mutant optic lobes also showed a significantly increased slope (Fig. 4I), which was measured from the coronal perspective (Fig. 4A). While the molecular mechanisms underlying these phenotypes will require further investigation and are beyond the scope of this work, these data nonetheless highlight the utility of using AI models to identify new mutant phenotypes even in animals that weren’t included in the training dataset.

In summary, our deep learning model facilitates fast, accurate, and consistent segmentation of the *Drosophila* brain with over 98% accuracy under a variety of Micro-CT imaging conditions. This allows for rapid characterization of brain mutant phenotypes and can serve as a tool for rapid quantitative analysis of potentially significant mutants. Our AI models reduce the burden of manual brain segmentation from ∼30 minutes to ∼3-5 minutes per brain, greatly enhancing the speed of our workflows. Key factors that determine the accuracy of a given model include image pixel size, image calibration and orientation, and consistency of staining procedures/methods. We note that Dragonfly’s AI platform can be extended into other forms of imaging beyond Micro-CT and electron microscopy that rely heavily on segmentation, including light microscopy. Our model is freely available for use and modifications using Dragonfly’s cloud-based services and we are actively building additional models for other organs, with the goal of having a trained model for each tissue to extend the usefulness of Micro-CT imaging and analysis for various applications in the fly.

## Materials and Methods

### Fly Stocks

Animals were maintained at 25C on cornmeal-agar. *asp*^T25^/*asp*^Df^ mutants were obtained by crossing *asp*^t25^/Tm6b and *asp*^Df^/Tm6b lines and selecting for the microcephalic phenotype (Schoborg et al., 2015). *Yellow white* (*yw,* Bloomington #1495) and *Oregon-R* (#25211) were used as wildtype. *eyD1Da* stocks were a generous gift from Patrick Callaerts (Callaerts et al., 2001).

### Microcomputed tomography

Staining and fixation: Flies were collected and stained following the protocol outlined here (Schoborg, 2020) for both Iodine and PTA stains. Imaging was performed on both the Zeiss Xradia Versa 610 and the Bruker Skyscan 1172, using the following imaging parameters:

### Skyscan

The source was set for 40 Kv, 250 uA, 10 Watts, with no filter. Images were taken using the medium pixel camera setting (2×2 binning), with a pixel size of 3.01 μm. The camera position was 80mm from the source and the object was 48mm from the source, with a 379 ms exposure time for iodine and 360 ms for PTA. The stage was set for a full 360-degree rotation, taking 900 projection images, using 3 frame averages and a random movement of 10. Reconstructions were performed using Skyscan’s nRecon reconstruction software.

### Versa

The source was set at 40 Kv and 3 Watts, with an LE1 filter and a 4x lens. Source distance was between 10.5-11 mm, and detector distance was set between 13.5-14 mm for a pixel size range of 2.95-3.01. Images were taken with an 850 ms exposure time for iodine stains, and a 1 s exposure time for PTA stains. 1601 projection images were taken with adaptive motion compensation. Reconstructions were performed using Zeiss’ automatic reconstruction tool.

### Deep learning and image analysis

Image analysis was performed using Dragonfly software (Comet Technologies Canada Inc) using the Deep Learning Module. A Dell Precision 7920 Tower workstation with an Intel Xeon Silver 4114 CPU, 2x Nvidia Quadro RTX 5000 Graphics cards, and 128GB of RAM was used for all rendering and Deep Learning.

The following criteria are essential for generating accurate models: 1) images must have the same pixel size; 2) consistent image orientation; 3) calibration of the image based on a consistent pixel intensity. Image pixel size can be changed using the Image Properties function. To consistently orient the images, we used the esophagus as a landmark for the deep learning field of view (Fig. 1A). We then derived a new image based on that orientation, followed by an extraction of the portion of the fly we intended to segment including a portion of air (background) and the surrounding pipet tip. The air and pipet tip is crucial, as this serves as the image pixel intensity calibrator. These have consistent pixel intensity values, unlike the sample itself or even the liquid surrounding the sample due to variability in sample staining and leaching. Once the image has been “prepped” using these steps, it can either be used to create training data or segmented using deep learning.

Training data for the models was generated by manual segmentation of images in the Deep Learning wizard. Approximately 20% of the data was manually segmented, or every ∼5^th^ frame. Training data was then exported and used in the Deep Learning tool. Model architecture was based off a Pretrained model from the Dragonfly Team with the following parameters: Depth level 7, initial filter count 32, input dimension 2.5D, input slices 3, Batch size 512, Epochs 100, patch size 64,64,1, stride ratio 0.25, Data augmentation factor of 10, horizontal and vertical flips, 180 degree rotation, zoom .9-1.1, shear 2.0.

### Time Savings over manual segmentation

Orienting the brain and deriving a new image takes 2-3 minutes for an experienced Dragonfly user. Deep Learning segmentation is 30-60 seconds using our workstation, and processing islands, generating meshes, and putting the data in excel is about a minute. This represents a 6-10-fold increase in segmentation speed compared to our manual segmentation analysis pipeline, which is ∼30 minutes per brain.

### Model Reproducibility

To determine the reproducibility of each model (Fig. 1F), this was achieved by training 10 different 1 or 3 brain models on the same brain(s) using identical training data for 50-100 epochs. Data reflects the comparison of the average volume of 10 brains by 10 replicates of the 1 and 3 brain models.

### Statistical Analysis

All statistical analyses and graph generation were performed using GraphPad Prism software (v. 10.3.1)

## Acknowledgements

We thank Sam Fay, Holden Bindl, and Lars Kotthoff for discussions and effort on an earlier phase of this project. We also thank members of the Schoborg lab for feedback. We also thank the WY INBRE Program for supporting the purchase of the Bruker SkyScan 1172 and the WY Science Initiative’s Center for Advanced Scientific Instrumentation (CASI) for providing access to the Zeiss Xradia Versa 610.

## Data Availability

All models can be downloaded from Dropbox here: https://www.dropbox.com/scl/fo/a66rjhu5sgs8jzhzj2399/AM34epNSJwTt5KW-C69IFs0?rlkey=lvm1aqizyltvsn247i4g7hkwr&st=hdvcf362&dl=0.

## Videos

**Video 1, related to Figure 1**: Comparison of manual segmentation vs deep learning segmentation for the 3 brain model. Yellow = Central Brain, Purple = Optic Lobes, Cyan = Lamina. Frames have been synchronized for the fly-through. Manual segmentation on the left, deep learning segmentation is on the right. Scale bar = 100 μm.

**Video 2, related to Figure 1**: Rotating 3D views of the manual segmentation (top) and the deep learning segmentation for the 3 brain model (bottom). Yellow = Central Brain, Purple = Optic Lobes, Cyan = Lamina.

**Video 3, related to Figure 4**: Rotating 3D views of a wildtype (top), *asp* mutant (middle), and *ey* mutant (bottom) segmented using the comprehensive model. Yellow = Central Brain, Purple = Optic Lobes, Cyan = Lamina.

